# A neural compass in the human brain during naturalistic virtual navigation

**DOI:** 10.1101/2024.04.18.590112

**Authors:** Zhengang Lu, Joshua B. Julian, Geoffrey K. Aguirre, Russell A. Epstein

## Abstract

A central component of wayfinding is the ability to maintain a consistent representation of one’s facing direction when moving about the world. In rodents, head direction cells are believed to support this “neural compass”, but identifying a similar mechanism in humans during dynamic naturalistic navigation has been challenging. To address this issue, we acquired fMRI data while participants freely navigated through a virtual reality city. Encoding model analyses revealed voxel clusters in retrosplenial complex and superior parietal lobule that exhibited reliable tuning as a function of facing direction. Crucially, these directional tunings were consistent across perceptually different versions of the city, spatially separated locations within the city, and motivationally distinct phases of the behavioral task. Analysis of the model weights indicated that these regions may represent facing direction relative to the principal axis of the environment. These findings reveal specific mechanisms in the human brain that allow us to maintain a sense of direction during naturalistic, dynamic navigation.

## INTRODUCTION

When navigating from place to place in the extended environment, humans and nonhuman animals keep track of spatial quantities that are useful for wayfinding, such as facing direction (heading), position, bearings to landmarks, and distances to local boundaries. Neurons responsive to these quantities have been identified in freely-moving rodents and have been extensively studied through electrophysiological recordings.^1–4^ This line of research is important for two reasons: first, because it reveals the neural correlates of an ecologically fundamental behavior; and second, because the brain systems involved in the coding of spatial information, such as the hippocampal formation and retrosplenial complex, are known to be broadly involved in memory and cognition. Indeed, it is widely believed that findings from spatial neuroscience will ultimately provide an avenue for grounding high-level cognition in well-defined neurocomputational mechanisms.^5^

For this promise to be fulfilled, however, it is necessary to draw strong connections between spatial codes observed in rodents and spatial codes observed in humans. This is challenging because it is not easy to identify the neural correlates of spatial navigation in humans. One approach is to record from neurons intracranially in presurgical epilepsy patients while they navigate through real or virtual environments. Studies using this method have provided much insight,^6–10^ but the method is limited by the fact that data collection is opportunistic, limited in temporal scope, and restricted to clinical populations. Another approach is to monitor brain activation with fMRI during virtual navigation. Studies using this method have identified brain systems that are engaged during navigational tasks^11–13^ and show activity that scales with univariate quantities such as distance to the navigational goal.^14,15^ However, this method is limited by the fact that the BOLD signal in fMRI has a sluggish response, making it difficult to assess spatial codes such as location or heading, which typically change on a faster timescale during dynamic navigation.

Previous fMRI studies have attempted to circumvent the limitations of the BOLD signal by breaking experimental sessions up into separate trials during which the spatial code of interest (e.g. location, facing direction, goal direction) does not change.^16–19^ This allows for the application of well-established adaptation or multivariate decoding methods for interrogating spatial codes. However, because these spatial codes are observed outside of a naturalistic navigational context, it is unclear how well they generalize to realistic navigational tasks. In the current study, we attempted to surmount these limitations of fMRI by collecting data in a naturalistic navigation paradigm, and then analyzing it using voxelwise encoding models.^20^ This analysis method has been previously used to examine representational codes in complex dynamic stimuli with temporally overlapping features, such as movies^21^ and audiobooks^22^. Thus, we thought it would be useful for examining spatial representations during dynamic navigation in a realistic environment.

We focus in particular on representations of Facing Direction (FD). This quantity, sometimes referred to as “heading”, is approximately equivalent to the head direction (HD) signal that is commonly studied in rodents. The ability to represent FD/HD provides an internal “compass” that is a central element of spatial navigation. An organism that knows which way it is facing is oriented in the external world, and thus can use internal representations of space (“cognitive maps”) to plan vectors of travel that will take it directly to its goal.^1,23^ It can also use its heading code to path integrate, thus updating its position vector as it travels, allowing it to form new cognitive maps in unfamiliar environments.^24^ Finally, an organism with an FD/HD code can mediate between egocentric and allocentric representations in memory, an operation that is believed to be essential for mental imagery, prospective thinking, and episodic recall.^25^ Thus, understanding how heading is represented is crucial for understanding how the brain mediates both spatial and nonspatial cognition.

In rodents, HD cells that can support a “neural compass” have been identified in many brain structures, including the presubiculum, anterior thalamic nucleus, and retrosplenial cortex.^26,27^ In humans, the existence of a similar mechanism is much less clear. Heading-modulated neurons have been identified in the medial temporal lobe, but their properties have not been investigated in detail.^28^ (See also ^8^). Neuroimaging studies using a trial-based approach have identified signals related to FD in several brain regions, including the medial parietal lobe (within the functionally-defined retrosplenial complex, or RSC), superior parietal lobe, entorhinal cortex (ERC), and thalamus.^17,28–37^ By design, these trial-based studies could only test differences between a small number of headings, and could not test heading codes within a dynamic, naturalistic context. One previous study used a voxelwise encoding model approach similar to the one employed here to identify heading-related codes during dynamic virtual navigation.^38^ However, the virtual environment in this study consisted of a single circular arena with a constant set of perceptual features, making it impossible to test whether the heading code was independent of the perceptual inputs or stable across different locations.

Here we go beyond this earlier work by examining spatial codes in participants performing a realistic “taxi-cab” task in a large-scale virtual city containing multiple subspaces and goal locations. To identify heading-related representations, we examined fMRI responses within individual voxels for two signatures indicative of FD codes: FD-related activity, and FD-related adaptation. The former is modulation of the BOLD signal as a function of FD at any point in time. The latter is a modulation of the BOLD signal as a function of the stability or change in FD. We then tested whether the FD-related activity codes exhibited key characteristics of a “neural compass” such as representing heading in a manner that it is stable across perceptual changes, task changes, and locations changes; and representing all possible headings, not just some. To anticipate, our results indicate that two regions of the brain—RSC and SPL—code for facing direction during dynamic virtual navigation in a manner that is indicative of an internal “compass”.

## RESULTS

### Participants performed a ‘taxi cab” task in two visually distinct versions of a virtual city

To examine neural responses related to spatial coding during realistic dynamic navigation, we scanned participants (N=15) with fMRI while they performed a taxi-cab task in two versions of a virtual city (city v1 and city v2; Fig. 1). The cities were spatially identical, with the same layout of roads, walls, and buildings, but they had different surface textures on the buildings and road to make them visually distinct. This design allowed the two versions of the cities to act as perceptual controls for each other when analyzing spatial codes. Participants alternated between searching for passengers and delivering them to stores that served as goal locations. Each passenger was delivered to 2 stores in sequence. The twelve stores were in the same locations in both cities.

**Figure 1.**
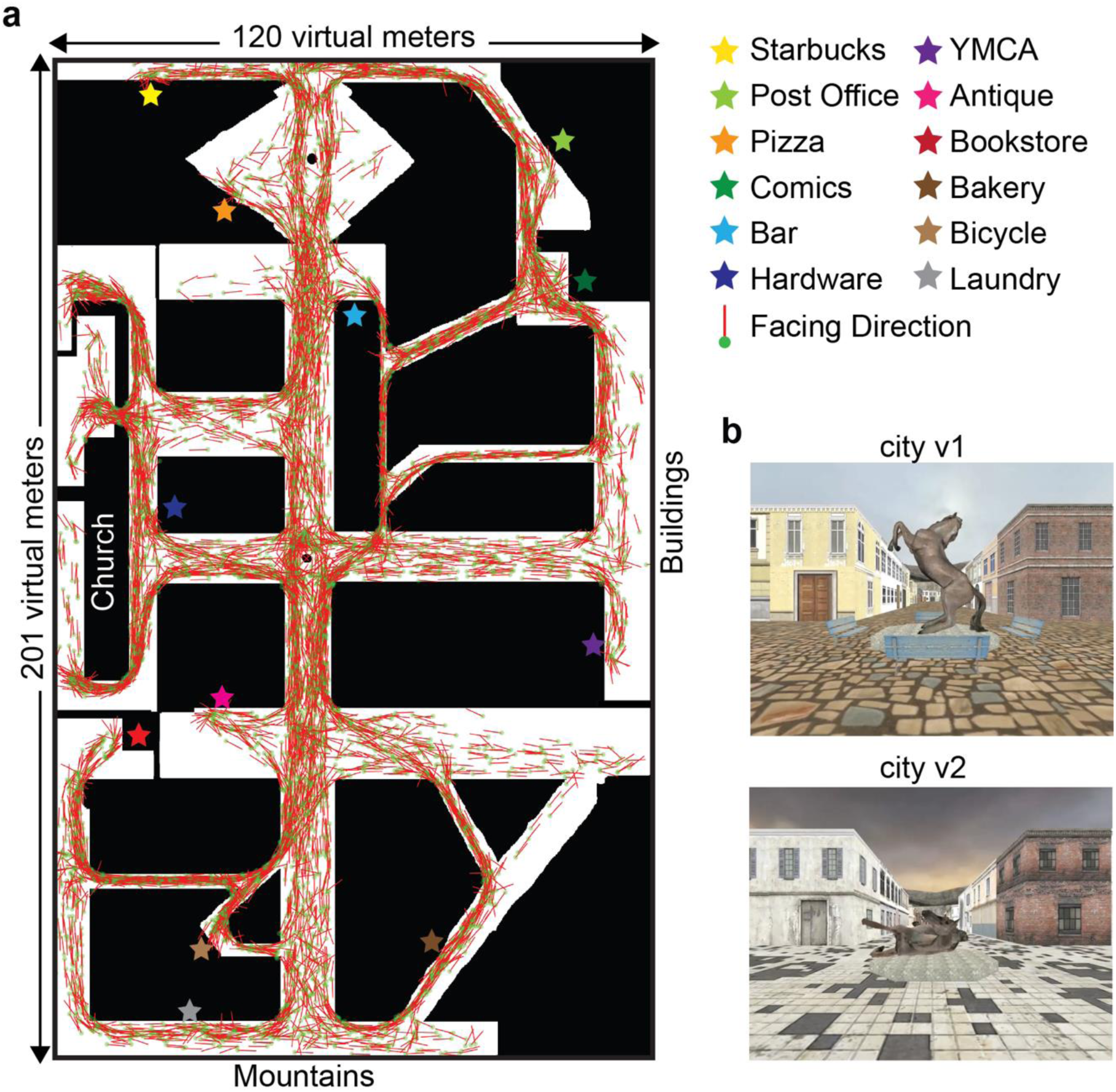
Active navigation in virtual reality (VR) cities. **(a)** Map shows the layout of the VR city; stars indicate the 12 target locations. The complete movement trajectory for one participant is plotted: green dots indicate the average location of the participant at each TR and red lines indicate the average facing direction. **(b)** Ground-level views showing the same location in the two versions of the city. The layout of the walls, buildings, and walkways is the same, but surface textures and colors are different. These two images show the initial view for each scan run, which was at center of the city facing to the “South” (i.e., towards the bottom of the map).

Participants were familiarized with the task and city v1 environment in two training sessions before beginning the scanning portion of the experiment in session 3 (see Methods). They were then scanned while playing the taxi-cab task in 5 or 6 different scan sessions spread out over a two-week period. The first three or four scan sessions were in city v1 while the remaining two sessions were in city v2 (See Supp Table 1). Each scan session was divided into multiple 11 min scan runs to avoid fatigue. On switching cities, participants were told that city v2 was a post- apocalyptic version of city v1, with the same spatial structure.

On average, participants found 4.08 passengers in each scan run, and delivered to 7.75 stores. The mean time to find a passenger was 98.73 s and the mean time to deliver to a store was 28.20 s. These values did not change over the course of the experiment (F(4,70) = 0.4845, p = 0.74, searching for passengers; F(4, 70) = 1.06, p = 0.3838, delivering to stores, Fig 2a), and did not significantly differ between city v1 and city v2 during store delivery (t(14) = 1.7793, p = 0.0969, Fig 2b) but did significantly differ during passenger search (t(14) = 2.1529, p = 0.0493; Fig. 2b). Visual examination of the trajectories taken by the participants indicated that they covered most of the environment when searching for passengers and took efficient routes when delivering passengers to their destinations (Fig. 1a). Examination of the histogram of facing directions indicated that all directions were sampled to some extent, but the cardinal directions (North, East, South, West) were sampled more, consistent with the rectilinear organization of the environment (Fig. 2c). FDs tended to remain constant over the course of a 2 s MR scan acquisition: on 67.03% of TRs, the FD remained within 8° (a magnitude corresponding to the bins of our analysis model; see below).

**Figure 2.**
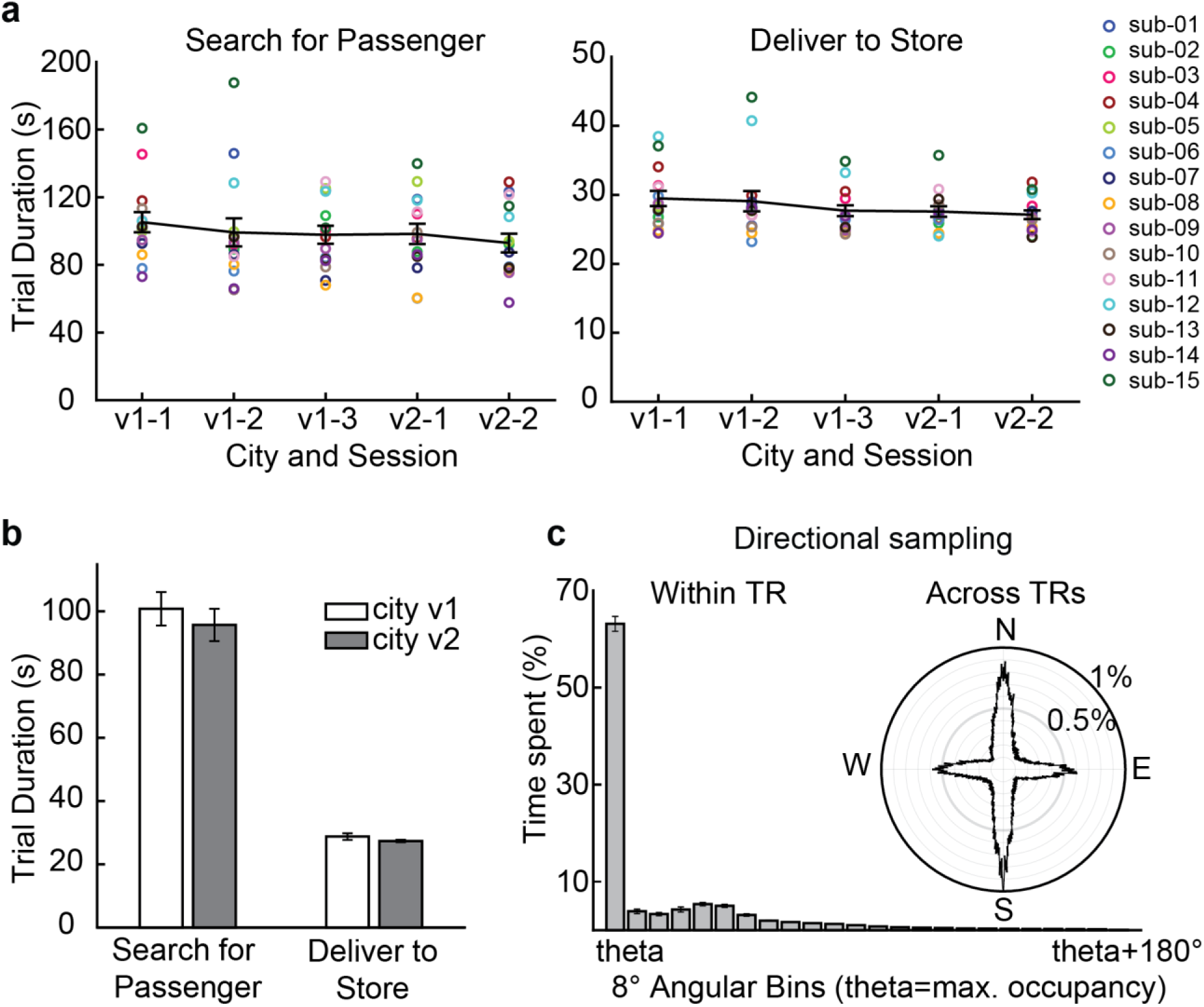
Navigation behavior. **(a)** Average duration of each trial phase, organized by scan session. **(b)** Average time of each trial phase, organized by city version. **(c)** Directional sampling during navigation. Within each TR, participants tended to face in a single direction for the entirety of that TR. Data were binned according to the angular distance to a predominant direction (theta) within each TR (8° bin). Across the entire experiment, participants faced all directions, but the cardinal directions (North, East, South, West) were sampled more, due to the rectilinear organization of the environment. Error bars indicate ±1 standard error of the mean.

### Motion energy-related signals were removed from the BOLD response

Our primary goal was to identify neural representations of facing direction (FD) by analysis of the fMRI BOLD signal. For any dynamic visual stimulus, motion energy will typically account for a large portion of the BOLD response. Visual motion energy is a particularly problematic confound when investigating FD codes, given the fact that changes in FD will invariably give rise to motion energy across the visual field. Thus, prior to analyzing the fMRI data for these FD-related signals, we removed responses related to visual motion energy.

To do this, we represented the motion energy of the visual display at every timepoint using a set of temporal-spatial Gabor filters. We then used ridge regression to fit the loadings on this motion energy feature space to the voxelwise BOLD response in each version of the city (city v1, city v2). Results for each city version were cross validated on the independent data from the other city version. Supplementary Fig. 1 shows that the motion energy model predicts BOLD signals across a wide swathe of cortex. We computed the residual between the predicted time course of the motion energy model and actual brain data for each city and z scored within each voxel across all time points for each scan run. The analyses of FD-adaptation and FD-activation reported below were performed on these residuals.

### fMRI adaptation for facing direction was observed in RSC and SPL

We next analyzed the fMRI response in each voxel for signals relating to FD-related adaptation; that is, modulation of the fMRI signal based on the stability of FD over time, under the specific hypothesis that the BOLD signal will be reduced if the same FD is maintained. These effects are of interest because regions that represent FD are likely to exhibit adaptation when the same FD is maintained over time. In addition, because FD is unequally sampled in our environment due to the rectilinear organization of the passageways, it is essential to account for FD-adaptation effects before considering FD-activation effects.

We modeled the FD-adaptation effect at each timepoint as the angular distance between the current FD and the integrated sum of preceding FDs. Preceding FDs contributed to the integrated sum with an exponential decay, such that more recent FDs had a stronger contribution. Because the parameters for the decay function (tau) were not known in advance, and were expected to vary across voxels, we fit the data using a nonlinear regression method that allowed us to search over these variables to find the best fit. Parameter search was performed on the data from each version of the city separately and then cross-validated on the independent data from the other version of the city.

Results are shown in Fig. 3. The two areas with the strongest FD-adaptation effect were in the medial parietal cortex, overlapping with the scene-selective retrosplenial complex (RSC), and the superior parietal lobe (SPL). Notably these regions have previously been implicated in the coding of information about facing direction (e.g. ^29,35^). We also observed FD-adaptation in the visual cortex, which was strongest in regions below the calcarine sulcus representing the upper visual field. This might be attributable to visual adaptation engendered by the stability of the visual input in the upper visual field when the navigator continues to face the same direction.

**Figure 3.**
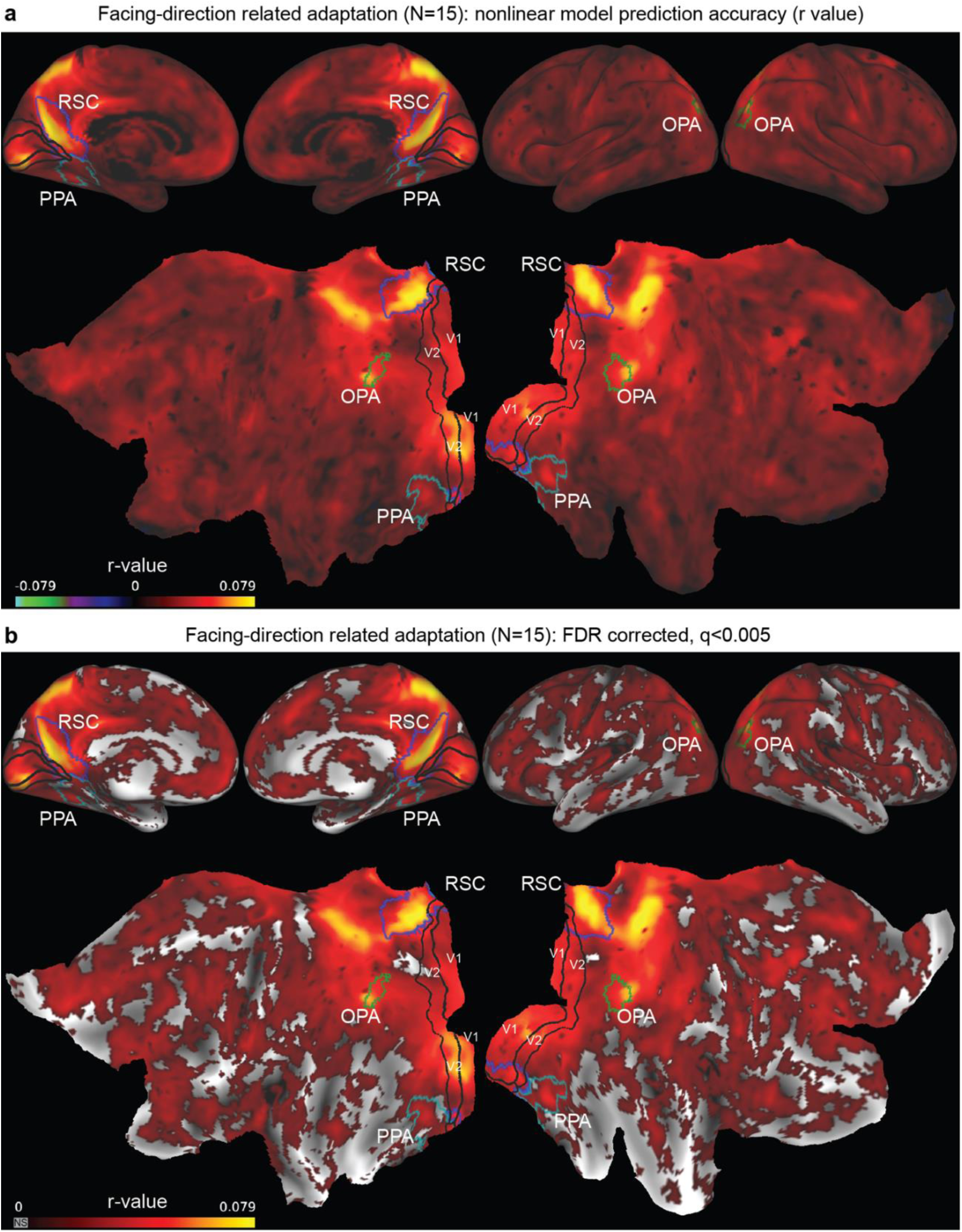
Facing-direction related adaptation. (a) Nonlinear fitting was used to estimate voxelwise parameter weights for an FD-adaptation model in city v1 and v2 data. Prediction accuracy was computed as the correlation (r) between the responses predicted by the model in one city version and BOLD activity in the other city version (cross-city validation). These values were then averaged across the two city versions and across all participants and plotted on a partially inflated brain (top) and flattened cortical surface (bottom). **(b)** Thresholded version of a (q > 0.005, FDR corrected). RSC=retrosplenial complex, PPA=parahippocampal place area, OPA=Occipital Place Area, V1/V2=Primary and Secondary Visual Cortices.

### Facing direction signals were observed in voxelwise fMRI activity in RSC and SPL

We then turned to an analysis of FD-related activity, which was our primary analysis of interest. To this end, we employed an encoding model approach in which the FD at each timepoint was represented as a response pattern across 45 circular gaussian filters (Supplementary Fig. 2). To account for potential FD-related adaptation effects that might be confounded with FD-related activity, we simultaneously modeled FD-adaptation using two regressors derived from PCA on the voxelwise adaptation effects obtained in the previously described nonlinear regression. We used banded ridge regression to fit these two feature sets (FD-activation, FD-adaptation) to the voxelwise BOLD response in each version of the city, and then cross-validated the results on the independent data from the other version of the city.

Results for the FD-activation model are shown in Fig. 4. There were two regions in each hemisphere exhibiting the strongest FD-activation effects: the medial parietal lobe, overlapping with RSC, and SPL. Results for the FD-adaptation model were very similar to those observed in the previously described nonlinear model (Supplementary Fig. 3).

**Figure 4.**
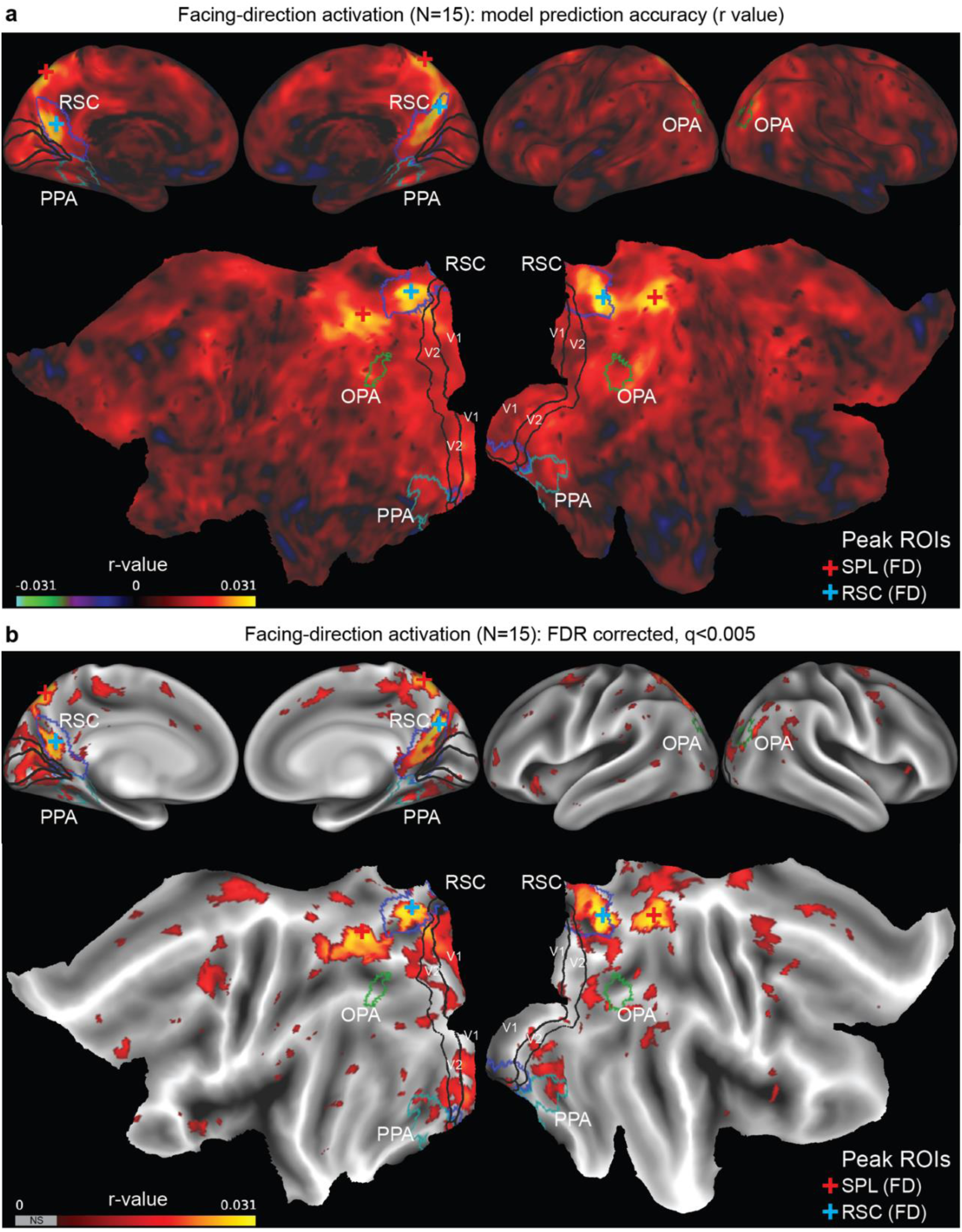
Facing-direction related activation. **(a)** Banded ridge regression was used to estimate voxelwise parameter weights for a combined FD-adaptation and FD-activation model in city v1 and v2 data. Prediction accuracy was computed for the FD-activation model as the correlation (r) between the responses predicted by the model in one city version and BOLD activity in the other city version (cross-city validation). These values were then averaged across the two city versions and across all participants and plotted on a partially inflated brain (top) and flattened cortical surface (bottom). **(b)** Thresholded version of a (q > 0.005, FDR corrected). ROIs same as Fig. 3. Crosses indicate peaks of the FD-activation effect in SPL (red) and RSC (blue), which were used to define SPL (FD) and RSC (FD) ROIs for further analyses (See 5c).

To test for consistency with previous findings, we also examined the FD-adaptation and FD- activation effects within six regions of interest that have previously been implicated in spatial processing during navigation (RSC, PPA, OPA, ERC, hippocampus, and thalamus), along with an early visual cortex (EVC) comparison region (Fig. 5). We found significant FD-adaptation in all seven regions (all ts(14) > 7.44, ps< 0.001, FDR corrected across ROIs) and significant FD- activation in RSC, PPA, OPA, ERC, thalamus and EVC (all ts(14) > 2.62, ps < 0.05, FDR corrected across ROIs) but not the hippocampus (t(14) = 1.665, p = 0.1182). Fig. 5c shows these same effects within RSC and SPL ROIs defined based on the RSC or SPL peaks in each participant showing the strongest FD-activation effect; these are used as ROIs in the analyses below.

**Figure 5.**
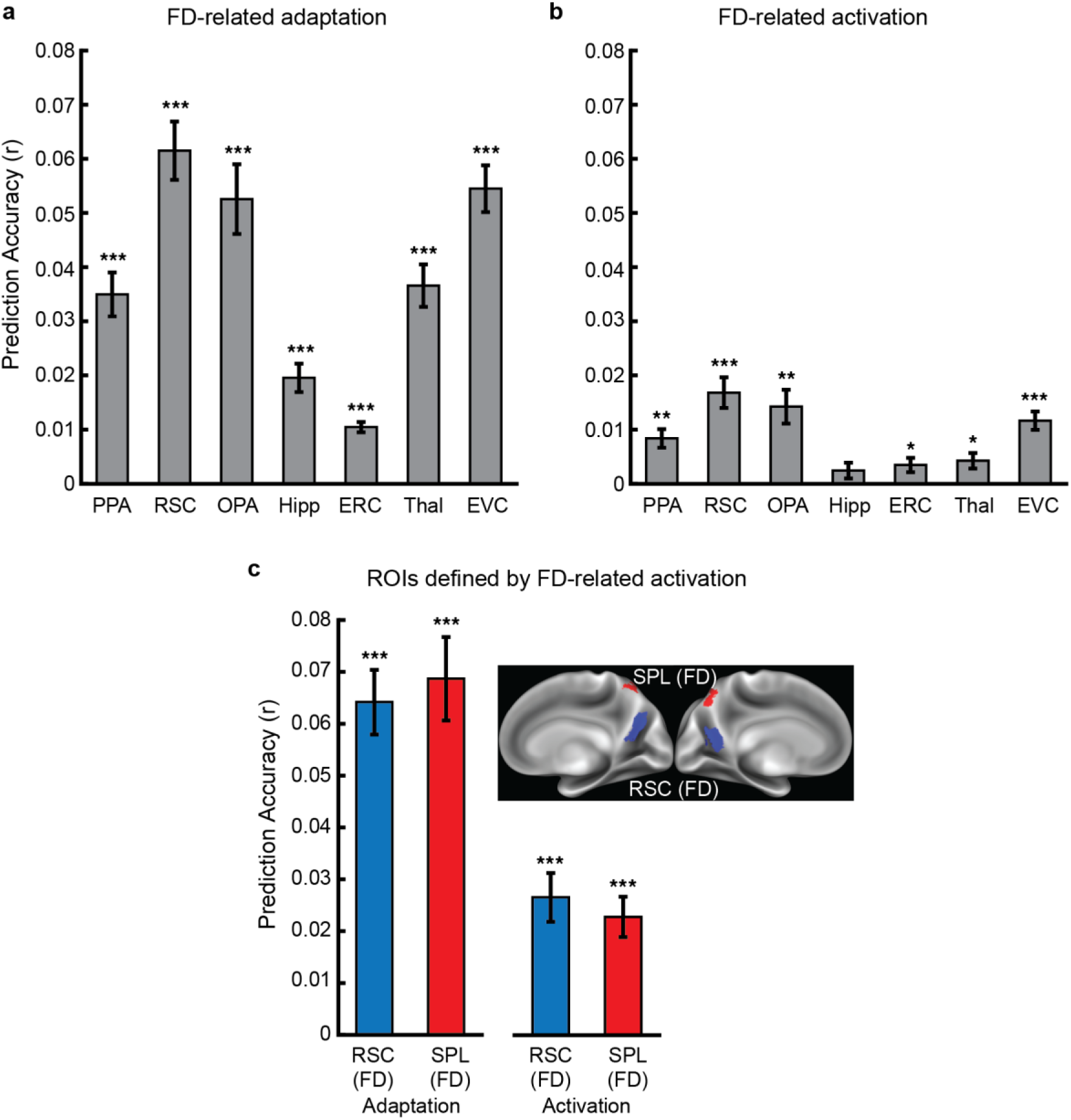
Facing-direction related coding in spatial system ROIs. **(a)** Significant FD-adaptation effects were found in all ROIs, including parahippocampal place area (PPA), retrosplenial complex (RSC), occipital place area (OPA), hippocampus (Hipp), Entorhinal Cortex (ERC), thalamus (Thal) and early visual cortex (EVC) **(b)** Significant FD-activation effects were found in all ROIs except hippocampus. **(c)** RSC (FD) and SPL (FD) ROIs were defined in each hemisphere as the ∼200 voxel showing the strongest FD-activation effect contiguous with the RSC (blue) or SPL (red) peaks. As expected, both FD-adaptation and FD-activation were highly significant in these regions. * p < 0.05, *** p < 0.001 (FDR corrected for multiple comparison across ROIs). Error bars indicate ±1 standard error of the mean.

### The direction code was stable across locations within the city and across phases of the experimental task

The above results suggest that RSC and SPL support representations of facing direction that could provide a “neural compass” during spatial navigation. For such a compass to be useful, it would need to be stable. The results above already provide evidence of stability across perceptual changes, given that the FD-activation model was trained and tested across different versions of the city that had the same spatial structure but different perceptual features. The results above also demonstrate stability over time, given that the city v1 and city v2 data were acquired in different scan sessions on different days. Here we test whether the FD-activation code also exhibits stability across locations within the environment and across different phases of the navigational task.

To examine stability across locations within the environment, we took advantage of the fact that our city was divisible into two subregions at a single choke point (Fig. 6a). We refer to these subregions as the “North” and “South” sectors of the environment. Using the same procedure as above, we trained the FD-activation model on each sector and tested it on data from the same or opposite sector in the other city. As before, we also included the FD-adaptation model in a banded ridge regression, and all analyses were performed on the residual data after regressing out BOLD signals related to motion energy. Fig. 6b shows results for the RSC and SPL regions that showed the largest FD-activation effect in the previous, whole-environment analysis (i.e., the ROIs shown in Fig 5c). The FD-activation effect was significant in both regions when analyzed within the same sector of the environment (RSC t(14) = 3.96, p = 0.0014; SPL t(14) = 5.64, p = 0.0001), and also—crucially—when analyzed across different sectors (RSC t(14) = 2.315, p = 0.0363; SPL t(14) = 3.0808, p= 0.0081). These results demonstrate that the FD-activation effect in these regions reflects a reliable directional code that generalizes across the entire environment.

**Figure 6.**
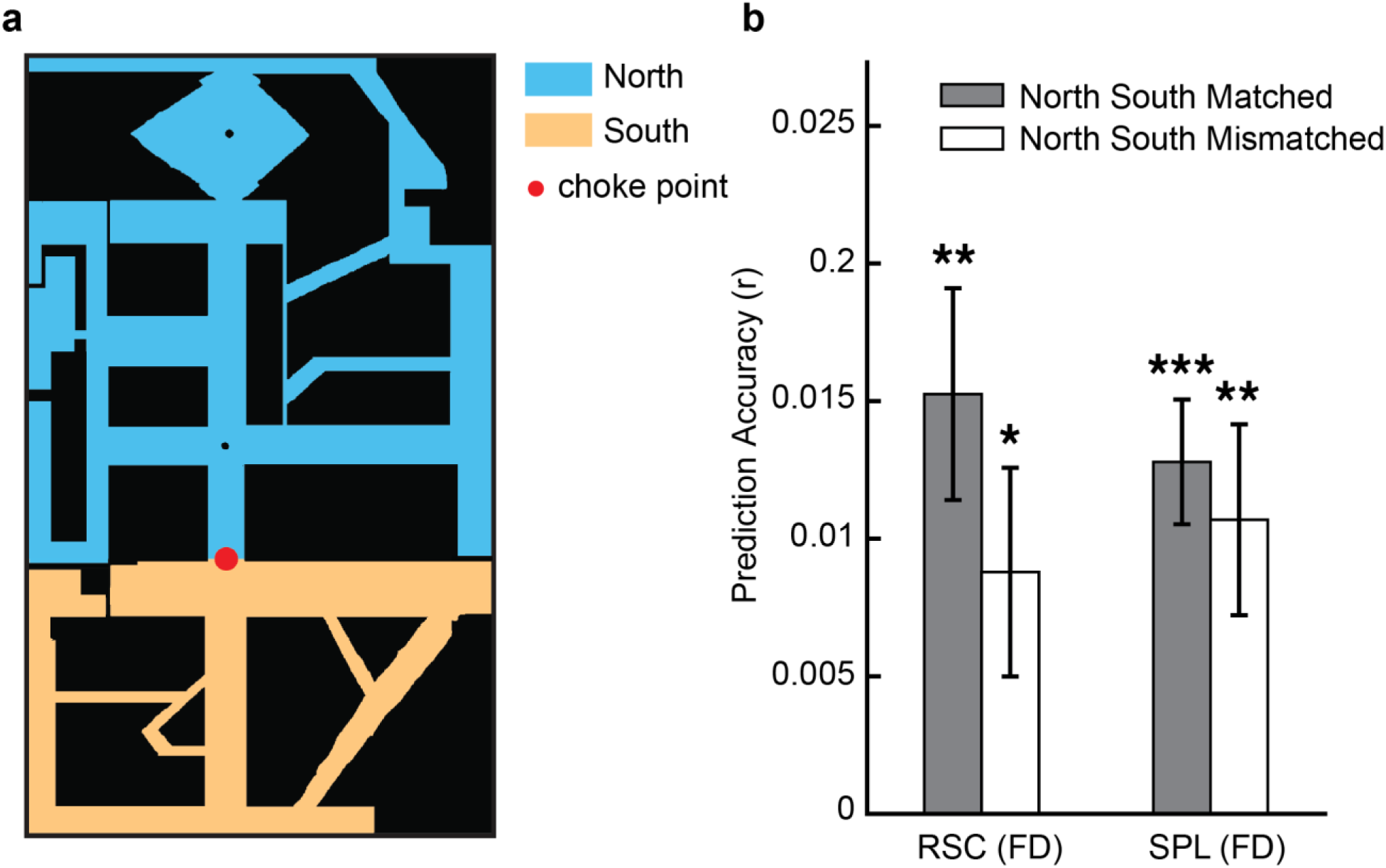
Facing direction representations are stable across locations in RSC and SPL. **(a)** A single choke point (red dot) divided the city into two sectors. The blue area indicates the “North” sector and the orange area indicates the “South” sector. **(b)** FD-activation models were trained on each sector separately and then tested on data from the same sector (gray bars) or opposite sector (white bars). Training and testing was performed across city versions (cross-city validation). Significant FD-activation effects were found for both matched and mismatched sectors, demonstrating that the FD code generalizes across locations. In RSC (FD) there was a reduction of the FD-activation effect for mismatched sectors, whereas no reduction was found in SPL(FD). * p < 0.05; ** p < 0.01; *** p < 0.001. Error bars indicate ±1 standard error of the mean.

To examine stability across different phases of the navigation task, we trained the FD-activation model separately on timepoints in which subjects were searching for a passenger and timepoints in which they were delivering a passenger to a destination. We then tested whether these two FD-activation models predicted BOLD activity in the corresponding or opposite task phase in the other city. Fig. 7 shows results for the RSC and SPL regions. The FD-activation effect was significant in both regions when analyzed within the same phase of the taxi-cab task, for both the search phase (RSC t(14) = 3.9943, p = 0.0013; SPL t(14) = 4.6534, p= 0.0004), and the deliver phase (RSC t(14) = 3.0397, p = 0.0088; SPL t(14) = 3.9838, p = 0.0014). Crucially, it was also significant when analyzed across the different phases of the task (search-deliver), (RSC t(14) = 4.4421, p = 0.0006; SPL t(14) = 2.7404, p = 0.0159). These results demonstrate the stability of the directional code across task phases with different motivational goals. Moreover, they indicate that the FD code is unlikely to be attributable to coding of goal (store) locations, since these are only relevant in the deliver phase.

**Figure 7.**
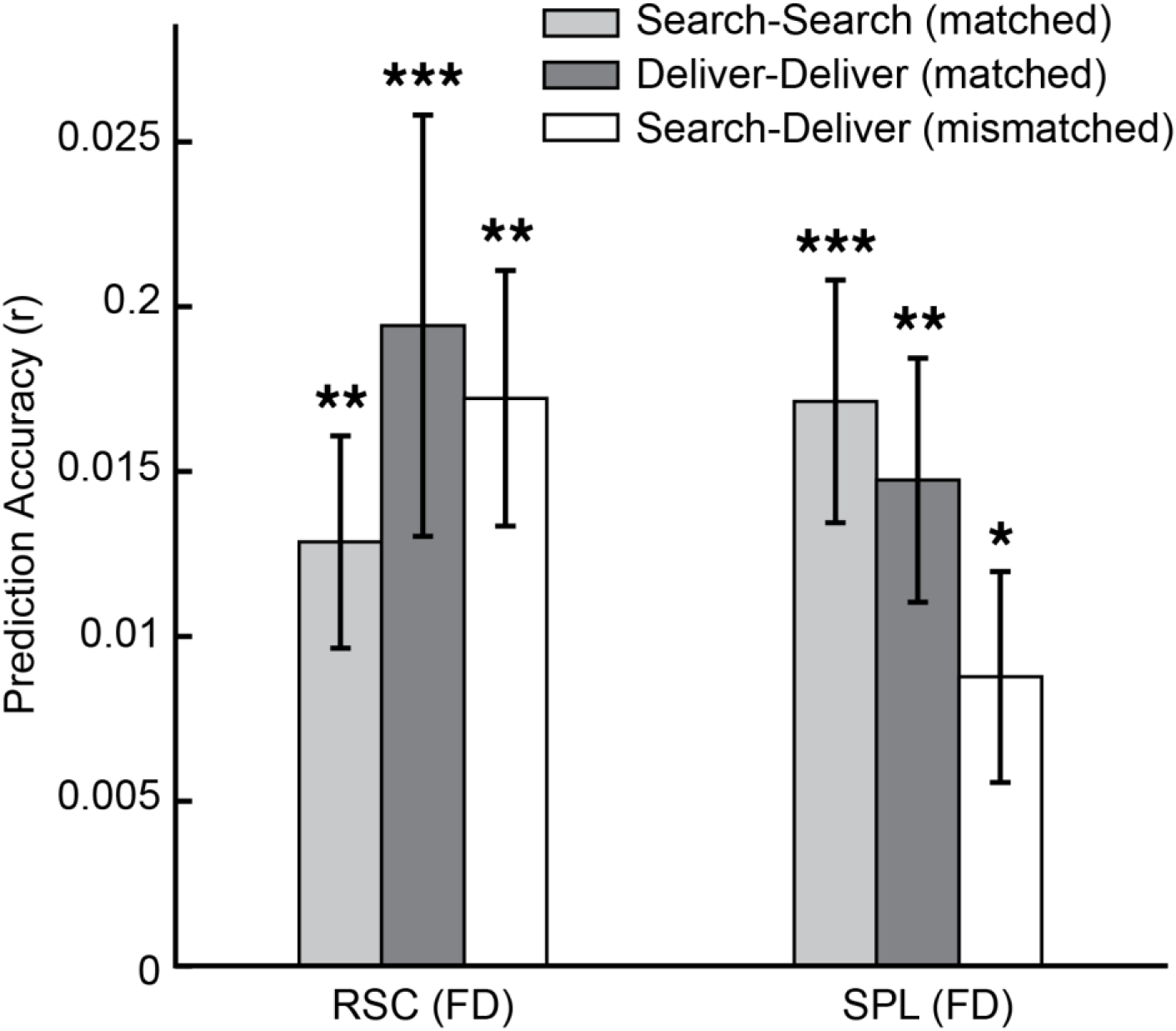
Facing direction representations are stable across task phases in RSC and SPL. FD-activation models were trained separately on search and deliver phases of the taxi cab task, then tested on data from the same phase or opposite phase. Training and testing were performed across city versions (cross- city validation). Significant FD-activation effects were observed for both matched and mismatched phases, demonstrating that the FD code generalizes across task phases with different behavioral goals. * p < 0.05; ** p < 0.01; *** p < 0.001. Error bars indicate ±1 standard error of the mean.

### Tuning profiles in RSC and SPL indicated that facing direction is coded relative to the principal axis of the environment

The analyses above indicate that RSC and SPL mediate FD codes that are stable across different versions of the virtual city, different sectors of the city, and different phases of the navigation task. We thus proceeded to investigate how FD is encoded in the estimated model weights in RSC and SPL. As a first step, we examined tuning profiles by plotting the parameter weights from each version of the city for two characteristic voxels (Supp. Fig. 4a) and the average parameter weights across all voxels within each ROI (Supp. Fig 4b). The individual voxels had parameter weights that varied as a function of FD, but there was no clear pattern in the average parameter weights. This is what one would expect if these regions represented the FD space: individual voxels should have FD tunings, but these tunings should differ across voxels and thus be eliminated by inter-voxel averaging.

We next examined the ensemble of voxelwise FD tunings. To do this, we normalized the weights for each voxel, and organized the voxels within RSC and SPL according to the angular direction of their peaks and troughs (Fig 8a). We saw an interesting and unexpected difference in the distribution of the peaks and troughs. The peaks were distributed across all possible FDs. This is an important finding, because it indicates that all possible directions are represented, as one would expect from a neural compass. The troughs, on the other hand, showed unequal distribution across FDs. Although all directions were represented to some extent, there was overrepresentation of “North” (270°) and “South” (90°)—corresponding to the principal axis of the city—in both RSC and SPL, as well as “East” (0°) in SPL. Analysis of the histograms for the peaks and the troughs (Fig 8b) shows clearly that troughs fall along these directions more often than would be expected by chance, whereas peaks are more equally distributed. This combined pattern of peaks and troughs demonstrates that the FD code in RSC and SPL is sensitive to the global structure (e.g., principal axis) of the environment. Indeed, it suggests the intriguing possibility that FD may be represented in these regions in terms of the angular deviation from the principal axis.

**Figure 8.**
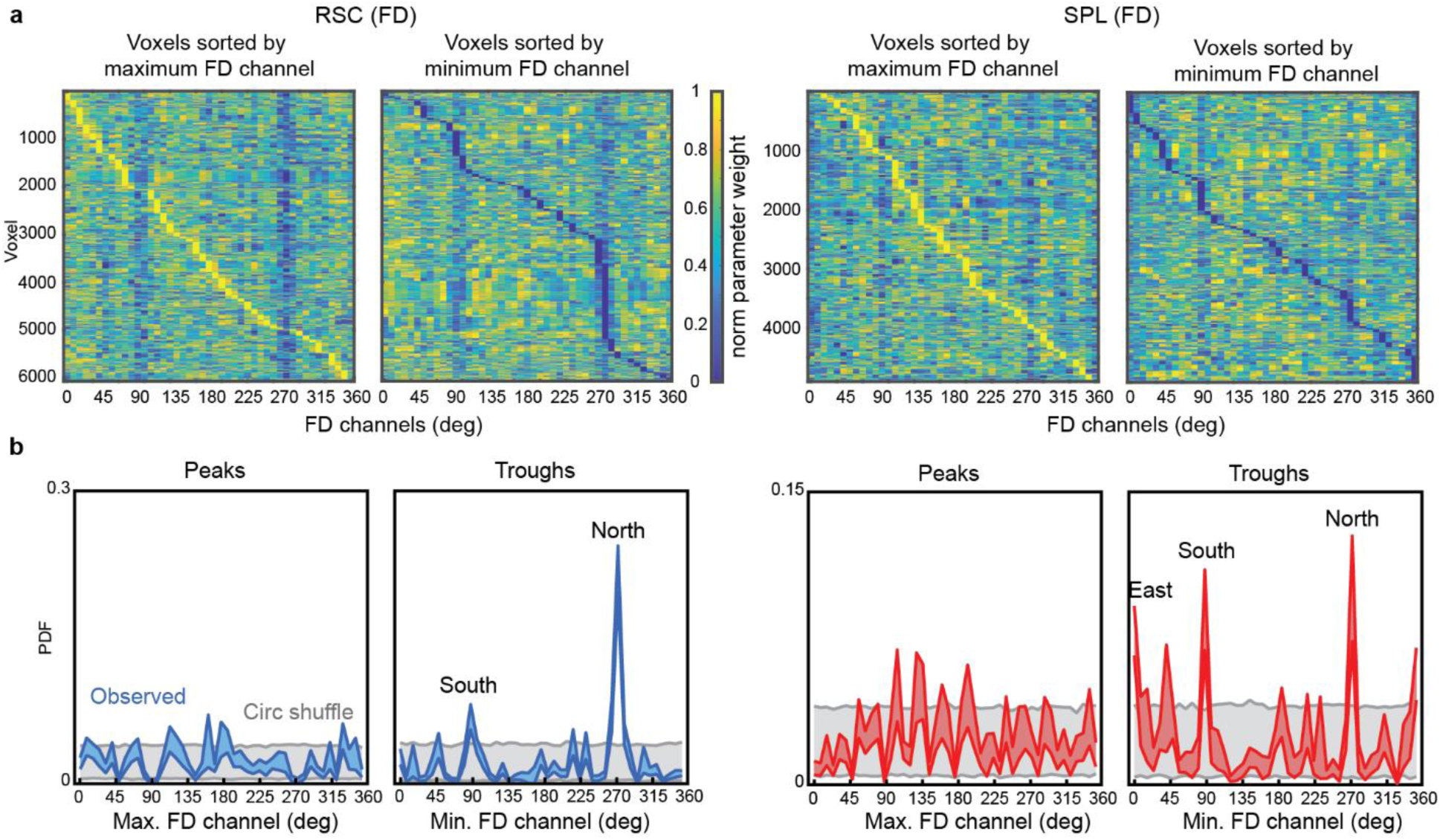
Analysis of model weights suggests coding of FD relative to principal axis of the environment. **(a)** Ensemble of FD-activation voxelwise model weights in RSC (left) and SPL (right). For each voxel, FD- activation model weights were averaged across city v1 and v2, and then normalized across the 45 FD channels. Voxels were then sorted according to the FD channel exhibiting the maximum (peak) or minimum (trough) parameter weight. Directions are equally represented in the peaks but unequally represented in the troughs. **(b)** Probability density function of maximum and minimum responses, constructed as a histogram of the voxelwise peaks and troughs. Peaks are evenly distributed across facing directions; troughs show over-representation of North and South in RSC, North South and East in SPL. Upper and lower colored curves are ±1 standard deviation of the observed data (bootstrap resampling). Gray range is ±1 standard deviation around chance baseline determined by circular shuffling of parameter weights across participants. Labelled peaks are significant outliers relative to the shuffled data (RSC: south p=0.03, north p<0.001; SPL: east p=0.01, south=0.006, north=0.003).

## DISCUSSION

The primary goal of this study was to identify FD codes in the human brain during naturalistic, dynamic navigation, comparable to the heading codes previously observed in freely moving rodents. To this end, we used fMRI to record brain activity while participants performed a realistic “taxi-cab” game in a virtual city over several experimental sessions. We identified two regions of the brain – RSC and SPL – that represented facing direction. These FD codes were evidenced in both adaptation (reduced fMRI activity when facing the same direction over time) and activation (voxelwise fMRI activity levels that varied as a function of facing direction). Crucially, the FD-related activation codes generalized across two different versions of the city, which had the same spatial layout but different visual appearances. They also generalized across spatially separated sectors of the city (North vs. South) and across temporally separated phases of the task (Search for passenger vs. deliver passenger to destination), indicating that they represent FD independent of location and behavioral goals. These findings emphasize the critical roles of RSC and SPL in representing directional signals during active spatial navigation. These regions may provide a neural “compass” that remains reliable as participants navigate from one part of the environment to another, akin to the function of head direction cells in rodents.

These findings build on previous studies examining FD (or “heading”) codes in the human brain but go beyond them in crucial ways. Several previous studies have used trial-based designs with adaptation or pattern classification methods to examine FD. In some of these studies, participants were given tasks that required them to implicitly or explicitly recall FDs from memory when cued by static verbal or visual stimuli.^18,28–31,35^ In others, they were shown movie clips depicting first-person movements through a virtual environment and asked to report the corresponding FD.^33,34^ These studies have found adaptation when FD is repeated on successive trials^28,31^, and multivoxel activation codes that distinguish between FDs on different trials.^17,18,29,30,32–35,37^ These effects have been observed in RSC and SPL, consistent with the present results, and also in ERC, thalamus, and subiculum. However, these previous studies did not examine FD codes while participants were actively navigating through an extended naturalistic environment, as we do here. Our findings show that the FD codes in RSC and SPL previously identified during constrained trial-based tasks are involved in representing facing direction during naturalistic navigation.

To interrogate spatial codes during dynamic navigation, we used a voxelwise encoding model. We represented FD at each timepoint as activity in a set of circular gaussian filters, each tuned to a specific direction, and we used regularized linear regression to relate these filter activations to BOLD responses in each voxel. This approach has been previously used to relate BOLD activity to visual, auditory, linguistic, and semantic features in other complex and realistic stimuli, such as visual movies and audiobooks.^21,22^ It was also used in a previous study to examine FD codes while participants dynamically navigated within a circular VR arena.^38^ This important precursor study found activation codes corresponding to FD in several ventral occipital and medial temporal lobe brain regions. FD responses were also observed in RSC, but—in contrast to the present results--RSC and SPL did not emerge as clear hotspots. The difference in anatomical distribution may be due, in part, to the fact that the previous study acquired fMRI data using a restricted set of fMRI slices that did not cover SPL and may not have covered all of RSC. In addition, it is possible that the RSC and SPL mechanisms we observed may be primarily engaged when participants perform a task that requires them to remain oriented within an extended space that includes several non-covisible subspaces.^39^ Such mechanisms might not have been as strongly engaged by the single-chamber environment used in the previous study. Indeed, because of the spatial restriction of the environment, the previous study could not test whether the heading code was stable across perceptual inputs and locations. In the current study, we show that FD codes in RSC and SPL generalize across perceptually different environments, locations within the environment, and task phases, thus providing stronger evidence that the observed responses relate to FD per se.

The use of an encoding model approach also allowed us to find important new information about *how* FD is represented in these regions. Specifically, when we examined the distribution of model weights across voxels, we found that the voxelwise peak responses were equally distributed across FDs. This is what one would expect from a “neural compass” in which all possible facing directions are represented. By contrast, the voxelwise minimum responses were not equally distributed, but were more commonly found for the “North” and “South” directions corresponding to the principal axis of the city. This finding is unlikely to relate to local geometry or boundaries, as these have a 4-fold symmetry related to the 90° offsets of the corridors along the four cardinal directions. Rather, this result suggests that FD is coded relative to the global layout of the city, which has a strong North-South organization along the main corridor. Notably, a recent study using a trial-based design found evidence that FD codes in RSC and SPL might be defined by angular displacement from a reference direction^35^, and we previously found evidence for preferential RSC coding of compass north when recalling imagined views from a familiar campus.^30^ Thus, we interpret our data as evidence that RSC and SPL represent FD as deviations from a reference direction, which in this case is defined by the global structure of the environment.

Although we observed the strongest FD effects in RSC and SPL, we also observed significant effects in several other ROIs, including ERC, PPA, OPA, and EVC, as well in other voxels throughout the brain. The finding of FD codes in ERC is consistent with previous literature using trial-based designs; however, it is notable that—though significant—the effects in this region were not particularly strong in the current experiment. This may suggest that FD representations in ERC are more strongly engaged in situations where FD must be newly recovered on each experimental trial, rather than situations when it must be maintained over a prolonged time period during continuous navigation. More broadly, a possible interpretation of our results is that FD codes are widely distributed across the brain, with some regions (RSC and SPL) being the primary but not exclusive nodes for this information. However, we note that this interpretation would be somewhat at odds with the neuropsychology literature, which suggests that deficits of the directional sense are related to lesions in circumscribed regions of the brain, specifically RSC, which is one of the two hotspots revealed in the current study.^40–43^

We also observed some evidence for differences between RSC and SPL spatial codes. Both regions represented FD in a manner that was consistent across spatially distinct sectors of the city (North vs. South) and temporally distinct phases of the taxi-cab task (search vs. deliver). However, cross-sector prediction accuracy was reduced in RSC compared to within-sector accuracy, whereas no similar reduction was observed in SPL. Conversely, cross-phase prediction accuracy was reduced in SPL compared to within-phase accuracy, whereas there was no similar reduction in RSC. These results indicate that FD codes in RSC are more impacted by changes in location, whereas the FD code in SPL are more impacted by changes in behavioral task.

One limitation of the current study—and all other previous fMRI studies of FD—is that we cannot separate FD codes related to the direction of the head from FD codes related to the direction of the trunk of the body. This is because the head and the body must remain stationary relative to each other during fMRI scanning. It is for this reason that we refer to the codes we observed as FD codes rather than HD codes, and we note that it is possible that these are different from the head direction codes observed in freely moving rodents. It is also the case that virtual navigation in the scanner is performed in the absences of vestibular or proprioceptive cues, which are known to have a strong influence on HD cells in rodents.^44^

These limitations aside, we believe that our approach holds promise for bridging the gap between the granular data obtainable from single-cell recordings in animals and the broad systems perspective provided by whole-brain fMRI in humans, as well as potentially revealing navigation mechanism that may be unique to humans and other primates. Although we focus here on FD codes, there is nothing intrinsic to the experimental design that prevents us from looking at other kinds of spatial codes. Indeed, a strength of the method is that one does need to set up distinct experimental paradigms targeted towards different spatial codes; instead, it is theoretically possible to examine multiple spatial codes that are simultaneously activated, and we expect to present such results in future reports. More broadly, the current study demonstrates an experimental approach that is likely to be widely useful for examining spatial codes during naturalistic dynamic navigation.

## MATERIALS AND METHODS

### Participants

Fifteen individuals (6 male; ages: 22-34, mean age: 27) were recruited from the University of Pennsylvania community and were paid for their participation. Written informed consent was obtained from each. All participants had normal or corrected-to-normal vision and reported that they were in good health with no history of neurological disease. Experimental procedures were approved by the University of Pennsylvania Internal Review Board.

### Experimental Procedure

#### Virtual cities

Two versions of the same virtual city (city v1 & v2) were developed using the Source video game engine SDK, Hammer Editor (valvesoftware.com). Cities v1 and v2 were spatially identical, with the same layout of roads, walls, and buildings, but they had different surface textures on the buildings and roads to make them visually distinct. City v2 was described to the participants as a “post-apocalyptic” version of city v1.

The cities spanned 120 x 201 virtual meters, and contained main streets, corridors, alleys, buildings, courtyards, a church, and statues. The edges of the cities were bounded by walls. A mountain outside the boundary was visible on one side of the city and apartment buildings outside the boundary were visible on another side. Twelve stores within the cities served as goal locations. These were clearly indicated by distinct storefronts and signage, and they were identical and in the same locations in both city versions. The store names were Antique Store, Swan Books, Post Office, Bicycle Store, Hardware Store, Bakery, YMCA, Laundry, Smith’s Bar, Comic Den, Starbucks, and Luigi’s Pizza.

#### Behavioral training

Prior to scanning, participants completed two 1-hour behavioral training sessions on a laptop computer to familiarize them with city v1 and the experimental task. During these sessions, participants navigated through the city from a first-person point of view and controlled their movements using arrow buttons on the keyboard (with left arrow button for turning left, right arrow button for turning right, and up arrow button for moving forward). The first training session consisted of three stages. In stage 1, participants freely explored the city for 15 min. In stage 2, participants navigated between the 12 target stores in a random order. The name of a target store was presented at the bottom of the screen; participants navigated to the named store, and upon reaching it they were prompted to find another store. This stage of training was self-paced and terminated once all target stores were found. In stage 3, participants played a taxi-cab game. The game alternated between a search phase and a deliver phase. In the search phase, participants navigated through the virtual city until they found a passenger, who they picked up by walking to the passenger’s location. The name of a goal store then appeared at the bottom of the screen, indicating that participants were to deliver the passenger to that location. Upon reaching the indicated goal store, the name of a second goal store appeared, indicating that participants were to further deliver the passenger to the second goal. The cycle then began again with the participant searching for a new passenger. Passengers could appear in 24 possible locations that were distributed throughout the maze so that passenger locations appeared to be random from the point of view of the participant. Participants played the taxi-cab game until the end of the 1-hour session. They were then asked to return at their earliest convenience for session 2 of behavioral training (14 out of 15 subjects came back within 2 days), during which they played the taxi-cab game on the laptop for an additional hour.

#### fMRI task

fMRI scanning took place over 5 or 6 separate scan sessions extending over a maximum of three weeks (mean = 13.4 days, see Suppl Table 1 for details), with the first session beginning 0-4 days after the second training session. In all scan sessions, participants played the same taxi-cab game that they performed in the behavioral training. Participants lay on their backs in the scanner and used a mirror mounted on the head coil to view a visual display presented at the end of the scanner bore on an InVivo SensaVue Flat Panel Screen at 1024 × 768 pixels resolution (diagonal = 80.0 cm, width × height = 69.7 × 39.2 cm). The visual display subtended ∼18.9° x 10.7° degrees. They controlled their movements with a 4-button MR-compatible button box (with left button for turning left, second button for moving forward, and third button for turning right). City v1 was used for the first three or four sessions and city v2 was used for the remaining two sessions (Supp Table 1). When switching cities, the participants were told “A war has broken out in your city. It is still the same city, but it looks different.” Each fMRI run was 11 min in length (330 TRs). Navigation on each run began with the participants standing in front of a horse statue at the center of the city and facing the South (Fig. 1b). The number of scan runs in each session varied slightly across participants depending on the time available (see Supp Table 1 for details). Total scan time in city v1 was 165-187 min (4950-5610 TRs) and total scan time in city v2 was 88-99 min (2640-2970 TRs). Immediately prior to entering the scanner on each scan session, participants performed 2 search-deliver cycles of the taxi-cab task outside the scanner to re-familiarize them with the task.

### MRI Acquisition and Preprocessing

Scanning was conducted at the Center for Functional Imaging at the University of Pennsylvania on a 3T Siemens Prisma scanner equipped with a 64-channel head coil. T1-weighted images for anatomical localization were acquired in each scan session using 3D magnetization-prepared rapid acquisition gradient-echo pulse sequence (MPRAGE) protocol (repetition time [TR], 2200 ms; echo time [TE], 4.67 ms; flip angle, 8°; voxel size, 1 × 1 × 1 mm; matrix size, 192 × 256 × 160 mm]). T2*-weighted functional images sensitive to blood oxygenation level-dependent (BOLD) signals were acquired using a multiband gradient-echo echoplanar pulse sequence (TR, 2000 ms; TE, 25 ms; flip angle, 70°; voxel size, 2 × 2 × 2 mm; multiband factor, 3; matrix size, 96 × 96 × 81). Field mapping was performed after each MPRAGE scan with a dual-echo (echo time (TE) = 4.12, 6.58 ms) gradient echo sequence with pulse repetition time (TR) = 580 ms, flip angle (FA) = 45°, pixel bandwidth = 260, voxel size, 3.4 × 3.4 × 4.0 mm; matrix size, 220 × 220 × 208 mm.

Data from the first nine participants were preprocessed using fMRIPrep 1.2.6 (Esteban, Markiewicz, et al. (2018); Esteban, Blair, et al. (2018); RRID:SCR_016216), which is based on Nipype 1.1.7 (Gorgolewski et al. (2011); Gorgolewski et al. (2018); RRID:SCR_002502). Data from the other six participants were preprocessed using fMRIPrep 20.2.1 (Esteban, Markiewicz, et al. (2018); Esteban, Blair, et al. (2018); RRID:SCR_016216), which is based on Nipype 1.5.1 (Gorgolewski et al. (2011); Gorgolewski et al. (2018); RRID:SCR_002502). Note that the number of T1-weights (T1w) images and the number of BOLD runs varied slightly across participants (see Supp Table 1 for details). The parameters below are for one typical participant using fMRIPrep 1.2.6.

T1-weighted (T1w) images were corrected for intensity non-uniformity (INU) using N4BiasFieldCorrection (Tustison et al. 2010, ANTs 2.2.0). A T1w-reference map was computed after registration of 5 T1w images (after INU-correction) using mri_robust_template (FreeSurfer 6.0.1, Reuter, Rosas, and Fischl 2010). The T1w-reference was then skull-stripped using antsBrainExtraction.sh (ANTs 2.2.0), using OASIS as target template. Brain surfaces were reconstructed using recon-all (FreeSurfer 6.0.1, RRID:SCR_001847, Dale, Fischl, and Sereno 1999). Spatial normalization to the ICBM 152 Nonlinear Asymmetrical template version 2009c (Fonov et al. 2009, RRID:SCR_008796) was performed through nonlinear registration with antsRegistration (ANTs 2.2.0, RRID:SCR_004757, Avants et al. 2008), using brain-extracted versions of both T1w volume and template. Brain tissue segmentation of cerebrospinal fluid (CSF), white-matter (WM) and gray-matter (GM) was performed on the brain-extracted T1w using fast (FSL 5.0.9, RRID:SCR_002823, Zhang, Brady, and Smith 2001).

For each of the BOLD runs, a reference volume and its skull-stripped version were generated using a custom methodology of fMRIPrep. A deformation field to correct for susceptibility distortions was estimated based on a field map that was co-registered to the BOLD reference, using a custom workflow of fMRIPrep derived from D. Greve’s epidewarp.fsl script and further improvements of HCP Pipelines (Glasser et al. 2013). Based on the estimated susceptibility distortion, an unwarped BOLD reference was calculated for a more accurate co-registration with the anatomical reference. The BOLD reference was then co-registered to the T1w reference using bbregister (FreeSurfer) which implements boundary-based registration (Greve and Fischl 2009). Co-registration was configured with nine degrees of freedom to account for distortions remaining in the BOLD reference. Head-motion parameters with respect to the BOLD reference (transformation matrices, and six corresponding rotation and translation parameters) were estimated before any spatiotemporal filtering using mcflirt (FSL 5.0.9, Jenkinson et al. 2002). BOLD runs were slice-time corrected using 3dTshift from AFNI 20160207 (Cox and Hyde 1997, RRID:SCR_005927). The BOLD time-series (including slice-timing correction when applied) were resampled onto their original, native space by applying a single, composite transform to correct for head-motion and susceptibility distortions. Functional data were smoothed with a Gaussian kernel (6mm full width at half maximum) using 3dBlurToFWHM (AFNI). The six motion parameters were regressed out from the fMRIprep preprocessed functional data using ordinary least-squares regression (custom python scripts), and polynomial trends up to third order. Finally, functional data were z-scored within each run before model fitting.

### fMRI Analyses

#### Overview

To test for fMRI responses corresponding to FD codes, we fit the BOLD response in each voxel with three encoding models. The first model accounted for responses related to motion energy in the stimulus display. The second model accounted for responses related to FD-adaptation that are induced by changes in FD. The third model accounted for responses related to FD in each possible direction. Because we expected signals related to visual motion energy to be large and dominant, potentially masking the FD-related signals of interest, we fit the motion energy model to the data first. We then removed these signals before proceeding with the FD-related analyses. For the latter analyses, we first analyzed FD-adaptation effects alone using a nonlinear fitting approach. Then we analyzed FD-activation and FD-adaptation effects, together in a linear model to account for potential common variance.

#### Motion energy

To determine visual motion energy signals, we preprocessed the video information that was presented to the participants in each scan run. The game engine recorded the visual display at 15 Hz and we converted these game files to mp4 files. We then downsampled these videos from 1024x768 to 128x96 and added black bands of pixels to the top and bottom to make each frame a 128 x 128 square. These downsized video files were converted into 128 x 128 x 3 x 44220 (height x width x colors x frames) matrices in unit8 format using MATLAB and converted from RGB values to L*A*B colorspace. The luminance channel values were retained for the encoding model analysis and the color values were discarded.

Next we quantified the motion energy in the pre-processed videos using a set of temporal-spatial Gabor filters. Each filter was defined by one of the three temporal frequencies (cycles/time window, min tf = 1.33, max tf = 2.67), one of the five spatial frequencies (cycles/image, min sf = 1.5, max sf = 24), and one of the 8 orientations of the Gabor wavelets. Filters spanned a square grid that covered each frame of the video images. Two Gabor filters with quadratic phases (0 and 90) were defined for each location, orientation and spatial frequency, and the values from these two Gabor filters were squared and summed to produce one feature value for each quadratic pair of filters. The resulting motion energy feature space consisted of 2139 Gabor feature channels. Prior to analysis, the time-course of activation across these filters was downsampled from the visual display frame rate (67 Hz) to the fMRI sampling rate (0.5 Hz, TR = 2s) by computing the mean feature channel values within TR.

Finally, we used an L2-regularized (ridge) regression to estimate model weights that map motion energy feature space onto the BOLD fMRI data for each voxel in each participant. This analysis was performed separately for city v1 and city v2. Feature channels were normalized by z-scoring the activation values within each channel across all time points for each scan run. Five finite impulse response (FIR) predictors at lags of 0, 2, 4, 6, 8 s were used to estimate the hemodynamic response for each feature channel at each voxel. A leave-one-run-out cross validation procedure was used to choose regularization parameters (alphas). For each fold of the cross-validation, motion energy model weights were estimated using 20 possible alpha values (1-1000, logarithmically spaced) and then used to predict responses on the withheld scan run. For each voxel, we chose the regularization parameter that yielded the highest mean performance accuracy across folds, as assessed by Pearson’s correlation (r) between the predicted and actual fMRI responses. We then repeated the regression on the entire dataset for each city version (v1 or v2) using the best-fit regularization parameter. The residuals between the predicted time course and actual brain data were computed for each city and used for modelling of the FD- adaptation and FD-activation effects, described below.

#### Facing Direction Adaptation

By definition, adaptation effects are effects engendered by changes (or lack of changes) in the quantity of interest over time. To examine these effects, we used a non-linear model that accounted for variation in the BOLD fMRI signal in each voxel in terms of FD history preceding each timepoint. FD values (expressed as orientation in degrees) were recorded by the game engine at 5.127 Hertz. This vector was downsampled to the fMRI sampling rate (0.5 Hz) using the circular mean prior to model fitting.

The non-linear model was under the control of parameters that adjusted the temporal integration of FD values and the gain of the effect of this behavioral history upon neural response. At each time point the recent history of FD values was weighted by a decaying exponential function defined by a time constant (tau; bounded between 0.01 and 20s). The modeled adaptation neural response at a time point was given as the (circular) difference between the current FD value and the recent history of FD values. The neural adaptation effect underwent multiplicative scaling under the control of a gain parameter, and then this neural prediction was convolved by a canonical HRF. A non-linear search (MATLAB fmincon) was used to identify the parameters that minimized the L2 norm of the model fit the BOLD fMRI data at each voxel. The R2 value of the model fit at each voxel was retained for voxel selection and display.

We wished to account for the effects of adaptation in subsequent, linear models. To do so, we derived a linear subspace of observed FD-adaptation effects from the non-linear fitting results. This was achieved by retaining the model fit of adaptation effects in each voxel. Each time-series vector was normalized to have unit variance. Then, we applied eigendecomposition on the covariance matrix of the predicted responses in each participant. We selected the two principal components corresponding to the highest eigenvalues. Finally, we transformed the original predicted adaptation responses onto the new 2-dimensional subspace by multiplying it with the eigenvector matrix. This allowed us to reduce the dimensionality of the predicted voxel responses while preserving the most significant information of facing direction related adaptation. These vectors were then used as covariates in subsequent linear analyses, described below.

#### Facing Direction Activation

To model FD-activation in each voxel, we first converted the scalar FD value at each TR into an FD vector consisting of 45 separate FD channels. Each channel was defined by von Mises circular probability density function. The peaks of these functions were equally spaced around the 360° circle with a separation of 8°, and each function had a width of 8° defined as the full-width-at-half-maximum (FWHM). The value of each channel at each timepoint was the value of the scalar

FD at each timepoint passed through the channel’s von Mises function. Thus, at each timepoint, each channel had a value that scaled with the angular distance between the scalar FD and the peak of the channel’s von Mises function, with the highest activation for the channel that peaked closest to the scalar value. All feature channels were normalized by z-scoring the values within each feature channel across all time points within each run in the fMRI dataset (city v1 or city v2). Five finite impulse response (FIR) predictors at lags of 0, 2, 4, 6, 8 s were used to estimate the hemodynamic response for each feature channel at each voxel.

A joint model that includes the FD-activation feature space and the 2 PCs constituting the FD- adaptation feature space was estimated for each voxel in the two motion-energy residual fMRI datasets (city v1 and city v2), using a modified version of ridge regression called banded ridge regression (Nunez-Elizalde et al., 2019). This procedure effectively decorrelates the features of the component models (in this case, FD-activation and FD-adaptation), by their covariance and the regularization parameters. Each voxel was assigned two different regularization parameters: one for FD-activation and one for FD-related adaptation. To find the optimal pair of regularization parameters for each voxel, a range of 11 alpha values was explored using the same leave-one-run-out cross-validation procedure used to fit the motion energy model. This allowed the joint model to effectively estimate the model weights for FD-activation and FD-adaptation, independently across all voxels in the brain. The pair of regularization parameters that yielded the highest mean performance accuracy was selected for each voxel. These regularization parameters pairs were then used to estimate the model weights for each voxel. To validate the FD-activation and FD-adaptation models for each city version, we used the parameter weights for each model to predict responses in the other city version (cross-city validation) and calculated the Pearson’s correlation (r) between the predicted responses and the actual fMRI time course in the validation city-version dataset. The two folds cross city model performance accuracies were then averaged for each voxel in each participant. To determine the significance of model prediction accuracy, we applied FDR-correction to obtain the multiple comparison corrected p-value (p < .05) for each voxel.

#### Generalization of direction code across location and task

We used cross-sector validation to test whether the observed direction code generalized across different locations within the environment. The cities were designed such that a single choke point divided them into separate sectors, which we designated “North” and “South”. Typically, participants would spend a substantial portion of time in one sector before passing into the other, making them hemodynamically separable from each other. We thus grouped the fMRI timepoints for each city version based on the corresponding sectors, and re-analyzed the fMRI data accordingly, using the banded regression procedure described above. The model fits were assessed through cross-sector cross-city validations (4 folds: city v1 North vs. city v2 North; city v1 North vs. city v2 South; city1A South vs. city v2 North; and city v1 South vs. city v2 South), and performance for same-sector validations (North/North and South/South) was compared to performance for different-sector validations (North/South and South/North).

Similarly, we used cross-phase validation to test whether the directional code generalized across different phases of the task in which participants pursued different behavioral goals (searching for passenger vs. delivering passengers to stores). For this analysis, we grouped fMRI datapoints according to phase of the experiment (search vs. deliver), and re-analyzed the data accordingly. The model fits were assessed through cross-phase cross-city validations (4 folds: city v1 Search vs. city v2 Search; city v1 Search vs. city v2 Deliver; city v1 Deliver vs. city v2 Search; and city v1 Deliver vs. city v2 Deliver), and performance for the same-phase validations (Search/Search, Deliver/Deliver) was compared to performance for different-phase validations (Search/Deliver and Deliver/Search).

#### Analysis of Voxelwise Tuning

To better understand how FD information was represented in RSC and SPL, we examined the distribution of FD-activation model weights across the ensemble of voxels. For each voxel, FD- activation model weights were averaged across city v1 and v2, and then normalized across the 45 FD channels. For each ROI, voxels were pooled across participants and sorted according to the FD channel exhibiting the peak (or trough) parameter weight. To test whether the distribution of FD channels exhibiting the peak (or trough) parameter weights across voxels differed from what would be expected by chance, we performed bootstrap resampling (k=1000) of participants and cities (v1/v2). Chance baseline was determined by the same bootstrap analysis, but randomly shuffling parameter weights circularly across FD channels (k=1000). Since within-participant parameter weights tended to be correlated across voxels within an ROI, each voxel’s parameter weights were shuffled by the same angular offset within each participant. Statistical significance was assessed by computing the probability that the observed mean across bootstrap resamples differed from the circularly-shuffled bootstrap distribution, separately for each FD channel.

#### Regions of Interest

In addition to whole-brain analyses, we also performed analyses on several regions of interest. These included brain regions previously implicated in processing of visual or spatial information relevant to spatial navigation (hippocampus, entorhinal cortex, thalamus, parahippocampal place area, retrosplenial complex, occipital place area), along with early visual cortex as a comparison region. The PPA, OPA and RSC and EVC were defined based on visual category contrasts (scene>object for PPA, OPA, and RSC; scrambled object>intact object for EVC) obtained from a group of 42 functional localizer participants previously run in our lab. The ROI for the entorhinal cortex was defined from each participant’s anatomical parcellation as obtained from Freesurfer. Bilateral hippocampus and thalamus were anatomically defined using FSL’s automatic subcortical segmentation protocol (FIRST). We also defined RSC (FD) and SPL (FD) ROIs in each participant to explore the nature of the facing direction code in the regions where evidence for this code was the strongest. These were clusters of ∼200 voxels contiguous with the RSC or SPL peaks in each hemisphere showing the strongest FD-activation effect.

## Acknowledgements

Supported by NIH grants R01EY022350 and R21EY027047 to RAE.

## Author Contributions

Conceptualization, J.B.J. and R.A.E.; Methodology Z.L., J.B.J., G.K.A., R.A.E.; Software Z.L., J.J. and G.K.A.; Validation Z.L. and J.B.J.; Formal Analysis Z.L., J.B.J., G.K.A; Investigation Z.L. and J.B.J.; Resource Provision G.K.A.; Writing—Original Draft Z.L. and R.A.E; Writing—Review & Editing Z.L., J.B.J., G.K.A. and R.A.E.; Supervision R.A.E.; Funding Acquisition R.A.E.

---

**Supplementary Table 1.** Scanning schedule. Participants were scanned while playing the taxi-cab task in 5 or 6 different scan sessions spread out over a two-week period. The first 3 or 4 scan sessions were in city v1 while the remaining sessions were in city v2. Each scan session was divided into multiple 11 min scan runs.

| Subject | City v1 (N of runs) |  |  | City v2 (N of runs) |  |  |
| --- | --- | --- | --- | --- | --- | --- |
|  | ses-01 | ses-02 | ses-03 | ses-04 | ses-05 | ses-06 |
| 1 | 6 | 5 | 4 | 4 | 5 |  |
| 2 | 6 | 5 | 5 | 1 City v1<br>4 City v2 | 4 |  |
| 3 | 6 | 5 | 4 | 4 | 4 |  |
| 4 | 6 | 5 | 4 | 4 | 4 |  |
| 5 | 4 | 5 | 6 | 4 | 4 |  |
| 6 | 4 | 5 | 6 | 4 | 4 |  |
| 7 | 6 | 5 | 4 | 4 | 4 |  |
| 8 | 4 | 5 | 6 | 4 | 5 |  |
| 9 | 4 | 6 | 6 | 4 | 4 |  |
| 10 | 6 | 5 | 4 | 4 | 4 |  |
| 11 | 6 | 4 | 5 | 4 | 5 |  |
| 12 | 6 | 5 | 6 | 4 | 4 |  |
| 13 | 6 | 4 | 5 | 4 | 4 |  |
| 14 | 6 | 4 | 4 | 2 City v1 | 4 | 4 |
| 15 | 6 | 5 | 4 | 4 | 4 |  |

**Supplementary Figure 1.**
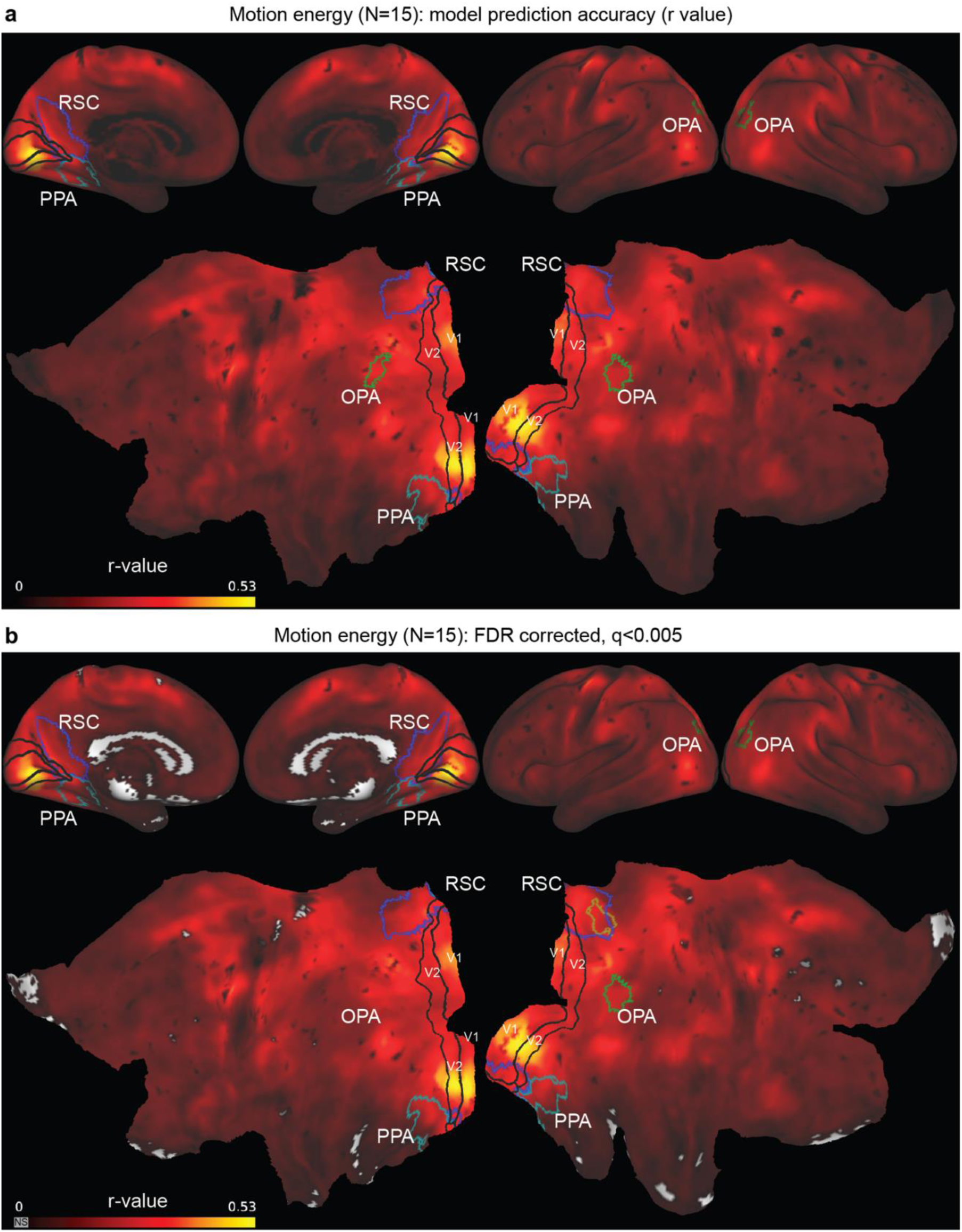
Motion energy related fMRI responses. **(a)** Ridge regression was used to estimate voxelwise parameter weights for a motion energy model in city v1 and v2 data. Prediction accuracy was computed as the correlation (r) between the responses predicted by the model in one city version and the BOLD activity in the other city version (cross-city validation) and plotted on a partially-inflated brain (top) and flattened cortical surface (bottom). Motion energy-related signals are found throughout the brain, with particular strength in early visual cortex. **(b)** Thresholded version of a (q > 0.005, FDR corrected). ROIs same as Fig. 3.

**Supplementary Figure 2.**
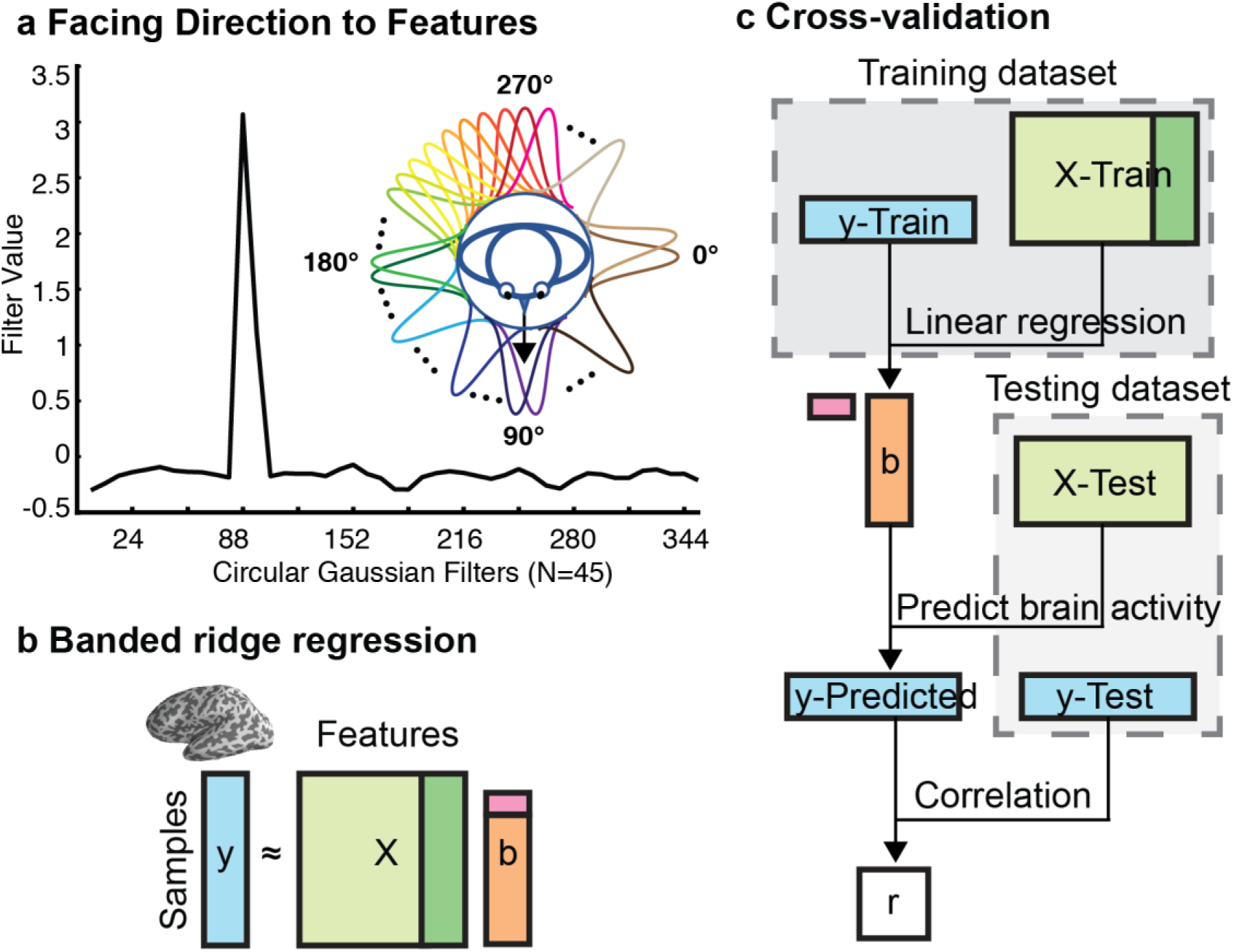
Voxelwise encoding model logic. **(a)** Facing direction (FD) activation feature space. We modeled facing direction using von Mises circular probability density function. At each TR, the observed FD was converted into a FD vector consisting of 45 separate channels. We generated a time course for all FD features. **(b)** A joint linear regression (banded ridge regression) was fit on two feature spaces simultaneously to predict brain activities (y). FD activation and FD adaptation feature spaces were concatenated into a large feature matrix (X). A weight vector b was estimated. L2-regularization parameters were estimated for each feature space separately in each voxel to improve the regression model. **(c)** Cross-validation procedure. We separated the data into a training dataset (e.g., city v1) and a test dataset (e.g., city v2). First, we estimated model weights (b) in each voxel in the training dataset only using banded ridge regression. We then used FD activation model weights on their own to predict each voxel’s brain activity in the testing dataset. Finally, we evaluated model prediction accuracy via Pearson correlation between the prediction vector and the recorded brain activity. Pearson’s R was used to quantify model prediction accuracy.

**Supplementary Figure 3.**
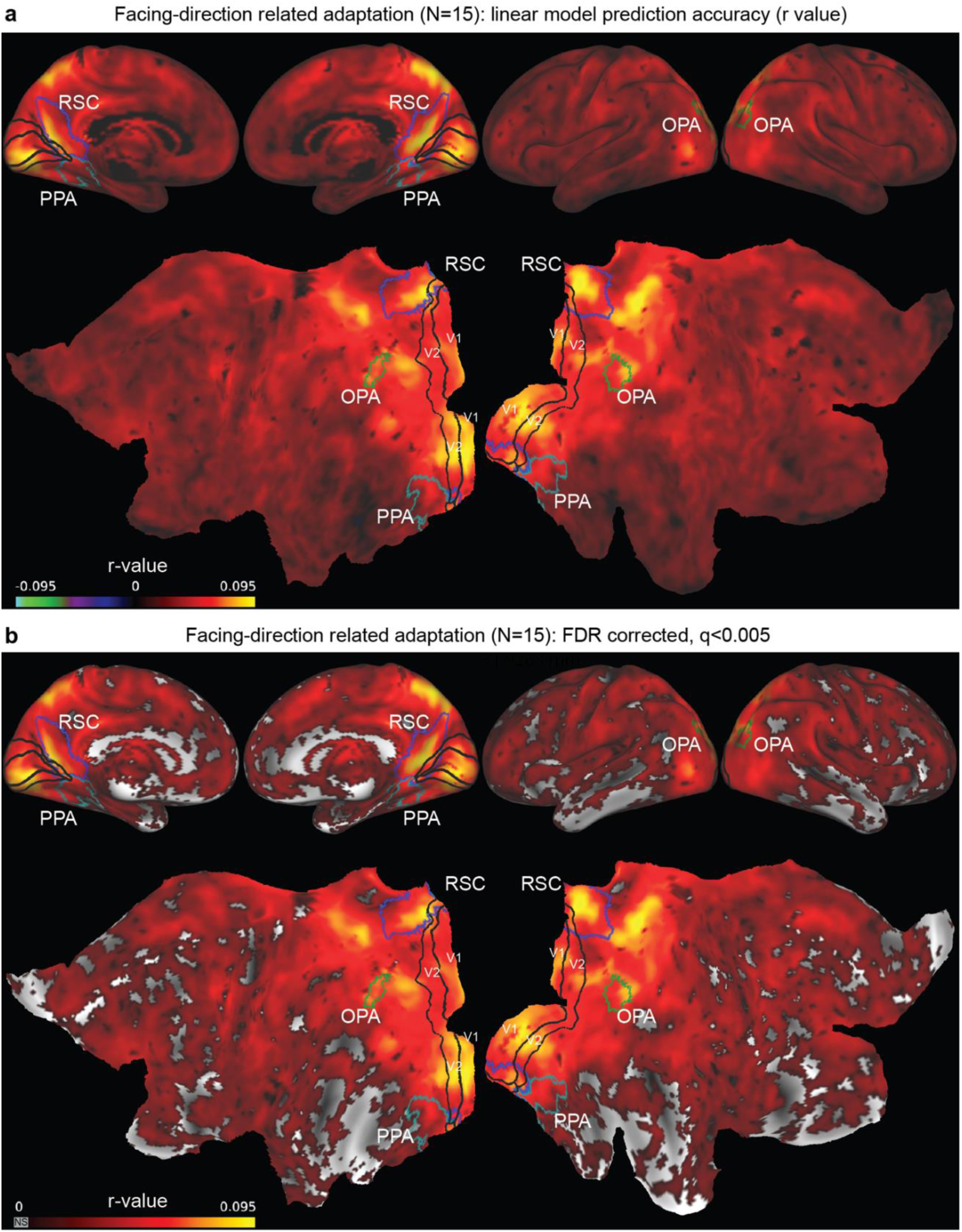
Facing-direction related adaptation in banded model. **(a)** Banded ridge regression was used to estimate voxelwise parameter weights for a combined FD-adaptation and FD-activation model in city v1 and v2 data. Prediction accuracy was computed for the FD-adaptation model as the correlation (r) between the responses predicted by the model in one city version and BOLD activity in the other city version (cross-city validation). These values were then averaged across the two city versions and across all participants and plotted on a partially-inflated brain (top) and flattened cortical surface (bottom). **(b)** Thresholded version of a (q > 0.005, FDR corrected). Note that the results are similar to those obtained with the nonlinear model (Fig. 3). ROIs same as Fig. 3.

**Supplementary Figure 4.**
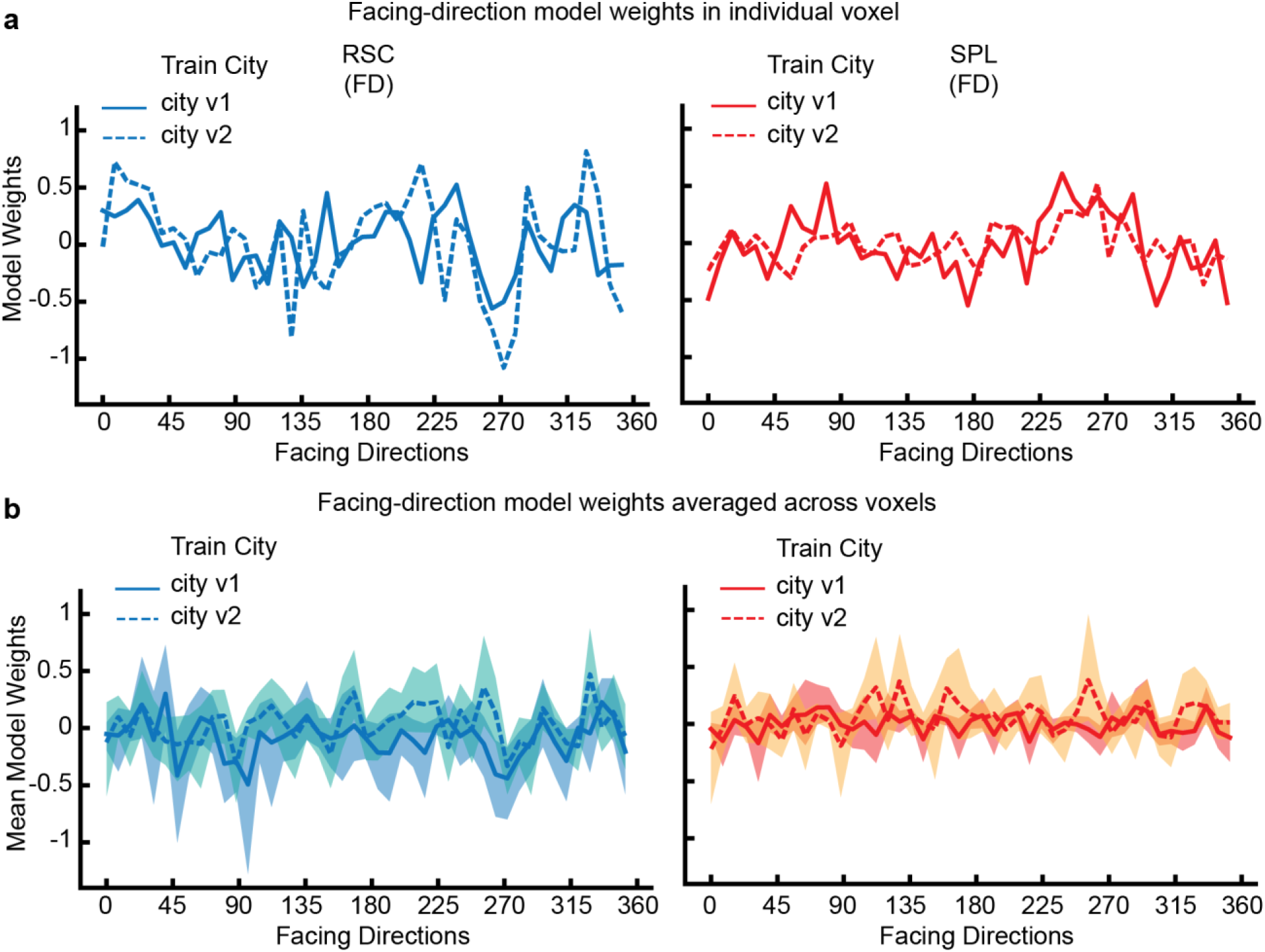
Model weights. **(a)** Model weights in single exemplary voxels. **(b)** Average model weights across all voxels in RSC (FD) and SPL (FD).

